# Panorama: a robust pangenome-based method for predicting and comparing biological systems across species

**DOI:** 10.64898/2025.12.22.695875

**Authors:** Jérôme Arnoux, Jean Mainguy, Laura Bry, Quentin Fernandez de Grado, Yazid Hoblos, David Vallenet, Alexandra Calteau

## Abstract

Over the last decade, the expansion in the number of available genomes has profoundly transformed the study of genetic diversity, evolution, and ecological adaptation in prokaryotes. However, traditional bioinformatic approaches based on the analysis of individual genomes are showing their limitations when faced with the sheer scale of the data. To overcome these constraints, the concept of pangenome has emerged, offering a comprehensive framework to capture the full genetic repertoire of a species. In this study, we present PANORAMA, an innovative pangenomic tool designed to exploit pangenome graphs and enable them to be annotated and compared in order to explore the genomic diversity of several species. Based on the PPanGGOLiN pangenome graphs, PANORAMA integrates advanced methods for rule-based prediction of macromolecular systems and comparative analysis of conserved features between different pangenomes, such as spots of insertion. We illustrate the use of PANORAMA on a dataset of 941 *Pseudomonas aeruginosa* genomes, evaluating its performance against reference defense system prediction tools such as PADLOC and DefenseFinder. The analysis was then extended to a larger set, including four species of Enterobacteriaceae (*>*6,000 genomes), demonstrating PANORAMA’s ability to annotate, compare, and explore the diversity and distribution of biological systems across multiple species. This work provides new methods for the large-scale comparative study of microbial genomes and underlines the relevance of pangenome approaches in deciphering their evolutionary dynamics. PANORAMA is freely available and accessible through: https://github.com/labgem/PANORAMA

**Author summary:** Microorganisms are present in nearly all environments on Earth. Uncovering their diversity through the study of their genomes is essential for understanding their biology and evolution. This includes characterizing the species’ complete genetic repertoire, known as the pangenome. Such research also enables new applications in health, ecology, biotechnology, etc. Here, we present PANORAMA, a novel computational tool designed to predict macromolecular systems, such as defense mechanisms against phages, and to compare pangenome graphs across different species. By using rule-based models that combine gene function and genomic context, we can search for systems directly within pangenome graphs. This graph-based approach provides a global view of the functional content of entire species, moving beyond the analysis of individual genomes. It greatly facilitates the analysis of thousands of genomes by reducing the required computation time and directly integrating the results to identify shared and specific systems. Furthermore, PANORAMA’s comparative functionality enables the identification of conserved structures across species, such as shared spots of insertion, revealing common evolutionary mechanisms and functional modules. This work establishes a foundation for comparative pangenomics, offering an unprecedented framework to explore the adaptive potential and evolutionary dynamics of prokaryotes at scale.

## Introduction

The rapid expansion of bacterial genome sequencing over the past decade has provided unprecedented opportunities to study the genetic diversity, evolution, and ecological adaptation of microbial organisms [1, 2]. For a significant number of species, sequences of hundreds or even thousands of strains are now available. While this wealth of information offers immense potential for discovery, it also presents significant challenges, as traditional genome-centric approaches, which focus on individual genomes, are becoming increasingly inadequate for managing and interpreting such large-scale datasets. To address these limitations, the concept of pangenome has emerged as a powerful tool. It encompasses the entire gene repertoire of a species, including *core* genes present in all strains and *accessory* genes found in only a subset of them, and provides a holistic view of genetic diversity and evolution within a species [3]. Pangenomics has significantly transformed microbial genomics by providing a comprehensive framework for understanding genetic diversity and functional capabilities across microbial species [4]. This approach allows researchers to investigate not only the genome of a single strain but also the complete gene repertoire within a species or group of strains, the pangenome, thereby enhancing insights into microbial evolution and adaptation.

Owing to the small size of their genomes and the large number of sequences available, particularly for species of clinical interest, pangenomic analysis of microbial genomes has benefited from the early development of tools, facilitating pangenome analysis, offering visualization, comparison, and partitioning of genomic data [5]. Among these tools, PPanGGOLiN stands out for its unique approach to analyze pangenomes by partitioning them with a statistical algorithm [6]. PPanGGOLiN represents genomic data as a pangenome graph at the gene family level, with nodes representing homologous gene families and edges capturing their genetic contiguity, enabling the compression of information from thousands of genomes while preserving the chromosomal organization of genes. A statistical model is applied to partition gene families in *persistent* genome (*i.e.*, gene families conserved in nearly all genomes) and variable genome, which includes the *shell* and *cloud* components corresponding to intermediate- and low-frequency gene families, respectively. PPanGGOLiN includes additional methods for the identification of Regions of Genomic Plasticity (RGPs) and their spot of insertion (panRGP method) [7] and their fine description in conserved modules (panModule method) [8], which have demonstrated their utility in identifying genomic islands and provide helpful insights into the genomic adaptability and evolution of bacteria.

Despite these advances, a significant challenge remains: detecting and comparing complex macromolecular systems at the pangenome scale. In microbial genomes, the genes responsible for macromolecular systems are usually arranged in a highly structured manner, typically clustered into one or several operons composed of functionally related genes. These clusters encode coordinated systems that play essential roles in microbial life. Among them are secretion systems, which allow the transport of proteins and other molecules across membranes to interact with the environment or host organisms; defense systems, which protect the cell from foreign genetic elements; and metabolic pathways, which organize enzymatic reactions to efficiently produce, transform, or degrade biological molecules. Understanding the organization and diversity of these systems is key to decoding the functional capabilities of microbial genomes. Several tools have been developed to detect macromolecular systems at the genome scale, including MacSyFinder [9], PADLOC [10], and DefenseFinder [11], the latter two being specialized in identifying bacterial anti-phage immune systems. They are highly effective when applied to individual genomes but are not designed to support systematic comparisons across large genomic datasets, such as detecting systems directly at the pangenome level. Tools capable of functionally annotating a pangenome at the scale of thousands of genomes remain limited. While some approaches can incorporate external annotations as additional metadata, such as PPanGGOLiN [6], Panaroo [12] and PanTOOLs [13], they do not support direct analysis of pangenomes, relying instead on predictions generally made at the genome level. Moreover, no tools enable the comparison of functional annotations across pangenomes from multiple species.

Here, we introduce **PANORAMA**, a powerful computational tool designed to harness bacterial pangenome graphs from large genomic datasets and enable comparisons across species to explore genomic diversity. Built on the PPanGGOLiN software suite, PANORAMA incorporates advanced methods for reconstructing and analyzing pangenome graphs. It offers several key features, including the ability to compare genomic contexts between pangenomes and annotate macromolecular systems at the pangenome scale. Functional annotation of biological systems is performed directly on the graph structures using rule-based models, making it possible to map and analyze complex genomic features without relying on linear genome representations. To illustrate the versatility of our approach, we focused on the comparative analysis and annotation of bacterial defense systems. Bacteria have evolved a remarkably diverse array of defense mechanisms against phages and other mobile genetic elements. These range from well-characterized systems, such as restriction-modification (RM) systems [14] and CRISPR-Cas complexes [15], to more recently discovered and less understood systems like BREX [16], DISARM [17], and retron-based defense systems [18, 19]. To date, over 150 systems have been described, unveiling an unsuspected diversity of molecular mechanisms [20]. This diversity is not only taxonomically widespread but also highly dynamic; defense systems are often associated with mobile genetic elements or genomic islands and can vary extensively in composition, organization, and presence both within and between closely related species [11, 21]. Despite this complexity, large-scale comparative studies of bacterial defense systems are still scarce. Pangenome-level analyzes hold great promise for revealing patterns of co-occurrence, horizontal gene transfer, evolutionary innovation, and defense strategies that are specific to particular species or lineages [22]. In this study, we present the methodology behind PANORAMA, a graph-based pangenomic framework designed to address these challenges. We first demonstrate its application on a comprehensive dataset of *Pseudomonas aeruginosa* genomes and compare its performance to established genome-scale tools such as PADLOC and DefenseFinder. We then extend this analysis to a broader dataset comprising four Enterobacteriaceae species, showcasing PANORAMA’s capacity to annotate, compare, and explore the distribution and diversity of phage defense systems across species. Overall, this work provides a powerful new resource for the comparative study of bacterial genomes and highlights the value of pangenomic approaches in revealing the evolutionary dynamics and ecological significance of bacterial defense repertoires.

## Results and discussion

### Overview of PANORAMA

To predict macromolecular systems, PANORAMA employs rule-based models similar to those used in MacSyFinder [9]. However, instead of applying these models to individual genomes, PANORAMA operates on the pangenome graph structure of PPanGGOLiN [6]. The rules rely on the presence/absence of specific functions predicted from pangenome gene families, incorporating constraints on their genomic organization (*i.e.*, gene colocalization). Functional annotation of pangenome graph gene families is performed through alignments with HMM protein profiles [23] defined for each macromolecular system. The genomic contiguity of gene families potentially involved in a system is then assessed on the pangenome graph by applying transitive closure and edge filtering. At the end, the predicted systems consist of sets of colocalized gene families from the pangenome graph, supplemented with information on their classification within the *persistent*, *shell*, or *cloud* genome, as well as their association with RGPs, modules, and spots of insertion. Systems are also projected onto the genomes to determine their presence and gene content in each strain. An additional functionality of PANORAMA is its ability to compare pangenomes, identifying similar systems and spots of insertion across species. Based on a set of predicted spots or systems in several pangenomes, PANORAMA computes a similarity score for each pair of elements by detecting shared gene families. Then, it applies a community clustering algorithm to group similar systems or spots into clusters. More details about PANORAMA methods are provided in the Materials and Methods section.

PANORAMA is available as open-source software written in Python and is designed for easy installation and usage to facilitate broader adoption by the community (https://github.com/labgem/PANORAMA). Software commands are organized into two main workflows (Figure 1). The PanSystem workflow begins by annotating gene families of the pangenome graph using the specified HMM library, and then applies system prediction rules from the model repository. Gene family annotations can also be performed externally and provided to PANORAMA by the user in a Tab-Separated Values (TSV) file. The PanCompare workflow performs comparative analyzes of two or more pangenomes, including gene family clustering and system/spot comparisons. Both workflows generate textual outputs (TSV files), graph-based representations (in GEXF or GraphML formats, compatible with Gephi [24] or Cytoscape [25] software for visualization), and figures to summarize results. Functional annotations and predicted systems are saved in the pangenome’s HDF5 file, allowing for further analysis. Additional utilities are provided to automatically convert system models and HMM libraries into the PANORAMA format, with support for models from MacSyFinder [9], DefenseFinder [11], CasFinder [26], and PADLOC [10]. Models are stored in JavaScript Object Notation (JSON) format, with a flexible and easy-to-understand grammar, enabling users to customize or create new models.

**Fig 1.**
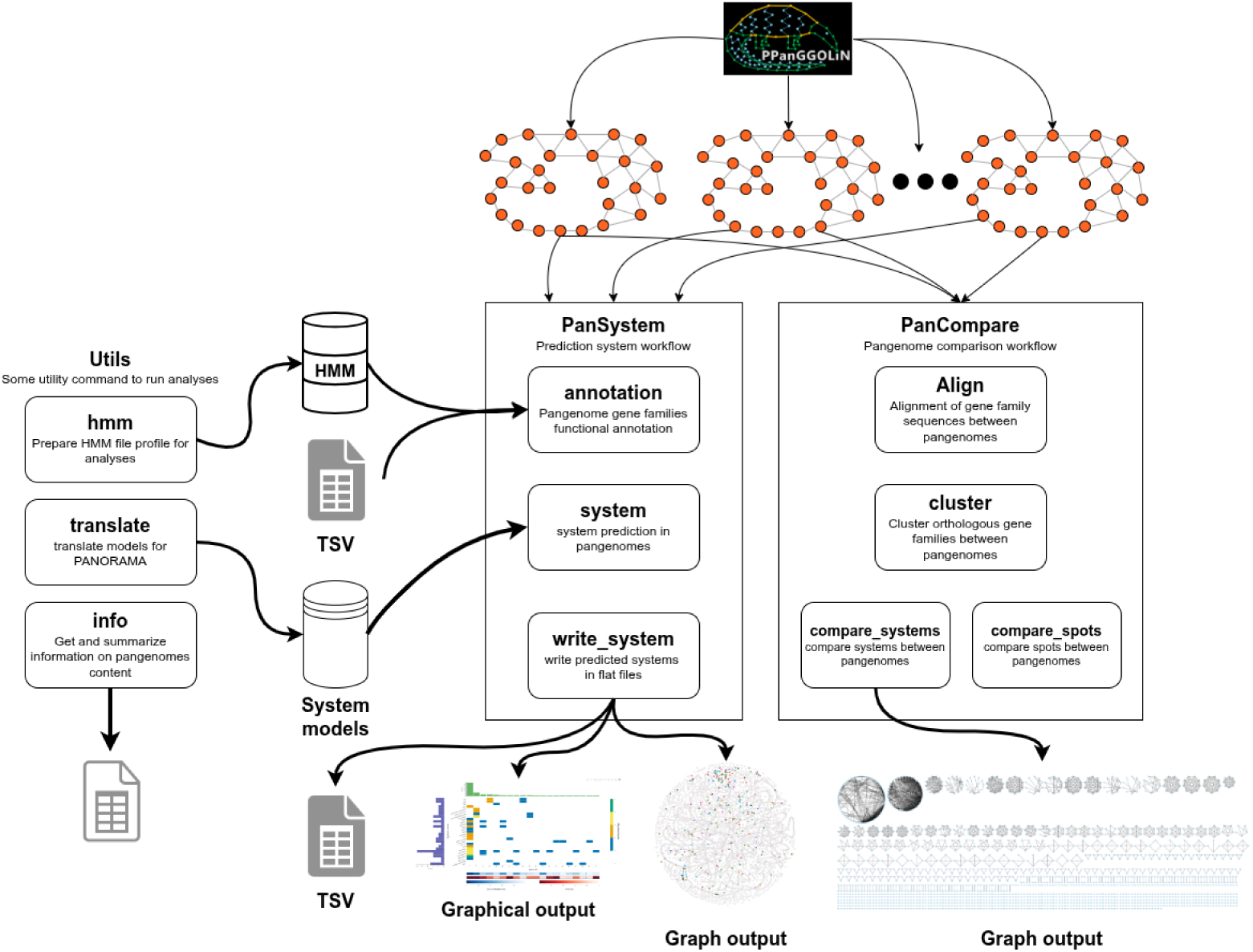
PANORAMA software overview. Each rounded box represents a possible software command integrated into workflows depicted in square boxes. PANORAMA is organized into two main workflows: *PanSystem*, which focuses on system prediction and annotation within pangenomes, and *PanCompare*, which handles comparative analyzes of pangenomes, including gene family alignment, clustering, and system/spot comparisons. Utility tools, such as HMM preparation and information summarization, facilitate data input and model translation. The software outputs include TSV files, graphical visualizations, and graph-based representations for comparing systems and pangenomes.

### System prediction benchmark

A set of 941 complete genomes of *Pseudomonas aeruginosa* was used to evaluate PANORAMA’s defense system predictions against two reference tools, DefenseFinder (including CasFinder models) and PADLOC. Although these tools use a similar approach to predict defense systems, they differ in the number of models (281 in DefenseFinder vs. 385 in PADLOC) and in the parameters used for predicting functions from HMM alignments, as well as for applying presence/absence and colocalization rules. Thanks to its generic system representation, PANORAMA is compatible with both tools and was run using their respective system models and HMMs after format conversion (*i.e.*, PanSystem workflow).

To conduct this benchmark, we assessed whether PANORAMA correctly assigned pangenome gene families to the appropriate systems based on the results from DefenseFinder or PADLOC. As expected, we obtained highly similar results, achieving an F1-score of 98.01% (recall: 98.84%, precision: 97.33%) using PADLOC as a reference and 96.64% (recall: 98.23%, precision: 95.10%) with DefenseFinder. As shown in Figure 2A, a substantial number of families are shared exclusively between PANORAMA and either DefenseFinder (650 families) or PADLOC (855 families), while only 964 families are common. This highlights the complementary nature of these tools and demonstrates that the pangenome analysis provided by PANORAMA is highly valuable for reconciling their results. Besides, PANORAMA missed only a few families: 56 for DefenseFinder and 47 for PADLOC (Table 1). These cases correspond to missing annotations in pangenome gene families. Unlike PADLOC and DefenseFinder, which analyze each sequence individually for every genome, PANORAMA relies on the protein sequence of the family’s representative gene for HMM alignments. Consequently, in rare instances, the alignment of the representative sequence falls just below the defined threshold, even though other sequences in the family have hits above it. Conversely, PANORAMA can identify additional results (318 families) for certain genomes whose gene sequence alignment with the HMM is less conserved than with the representative sequence. A last point concerns the systems predicted only by PANORAMA (*i.e.*, 136 with DefenseFinder models and 72 with those of PADLOC, see Table 1). Although such a high number was not initially expected, a detailed analysis revealed that some of these additional systems arise because PANORAMA’s genomic context search, based on colocalization rules in the pangenome graph, is less stringent than that of PADLOC and DefenseFinder. This relaxed approach allows PANORAMA to detect additional genes that are consistently conserved alongside known system components (see Materials and methods for further explanation). Other additional predictions stem from differences in HMM annotation due to the use of representative sequences for families, as mentioned earlier, but also from a bug in DefenseFinder that affected only RM and Retron systems, where alignment thresholds were not properly handled by the software. This bug has since been corrected by the authors (version 2.0.0) though it was not tested in this study. Other discrepancies arise because PANORAMA allows multiple associations between gene families and systems, whereas PADLOC and DefenseFinder link each gene to only one system. This is especially noticeable for systems with closely related models; for example, PANORAMA often predicts multiple Retron, RM, or Lamassu systems when other tools identify only one (Figure 2B). One potential improvement would be to include *forbidden* families in the models to facilitate their differentiation or to implement a scoring function, as done in MacSyFinder, to identify the best system. PANORAMA may also predict additional systems at the genome level for which the colocalization rule is not respected. These systems, flagged as *split*, have their genes separated, possibly due to rearrangements, insertion events, or assembly breaks occurring in a subset of genomes in which they are predicted.

**Fig 2.**
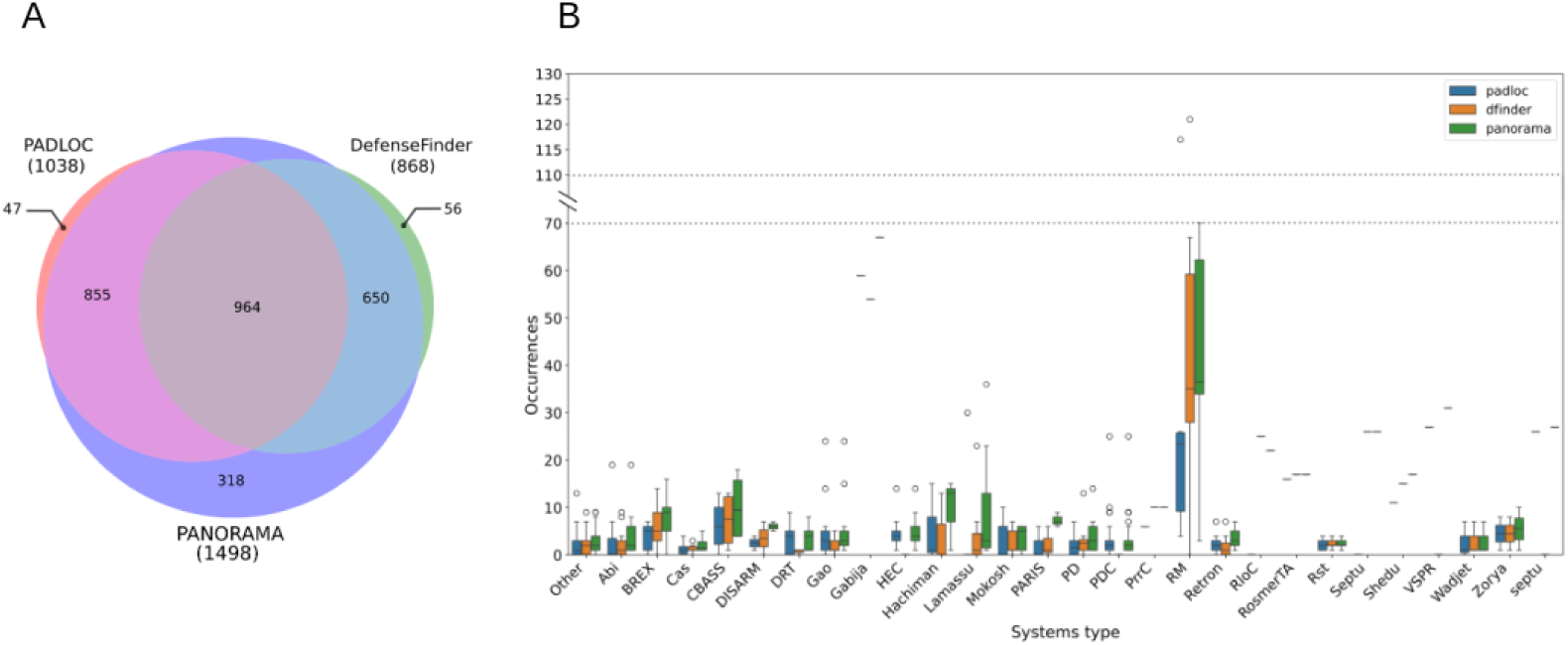
Comparison of PANORAMA, PADLOC, and DefenseFinder system predictions at the pangenome level. PADLOC and DefenseFinder predictions at the genome level were unified at the pangenome level by converting sets of system genes to sets of gene families. (A) Venn diagram illustrating the overlap of system families predicted by the three tools. (B) Boxplots displaying the distribution of system counts for each category, as predicted by the different tools.

**Table 1.**
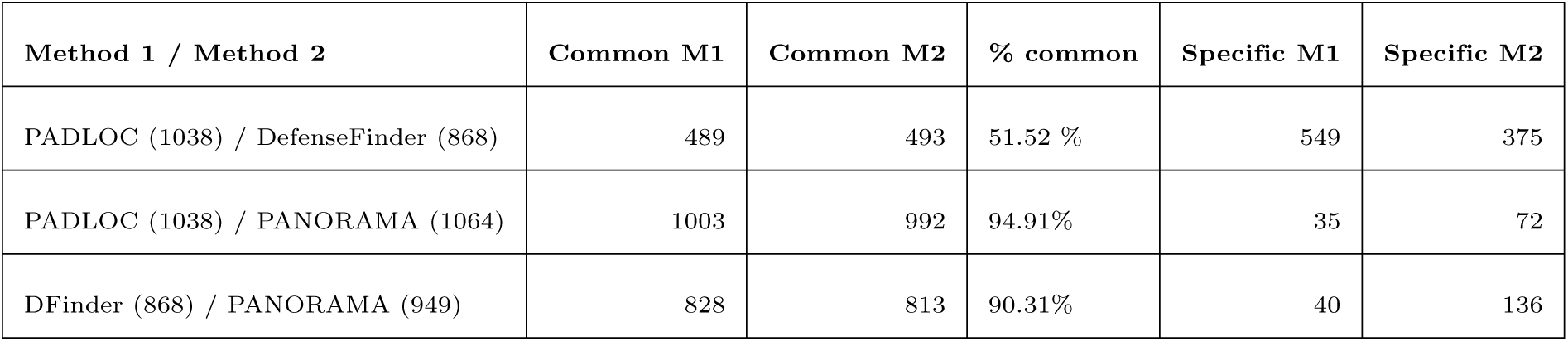
System prediction comparison between PANORAMA, PADLOC and DefenseFinder. For each comparison, M1 refers to the first method listed (Method 1) and M2 to the second method (Method 2). Common M1 and Common M2 indicate the number of systems predicted by both methods and attributed to M1 and M2 respectively. Specific M1 and Specific M2 represent systems uniquely predicted by each method. Numbers in parentheses indicate the total number of systems predicted by each method.

Tool performance execution was also evaluated by measuring their runtime and memory usage on a Linux server with 36 CPU cores (Table 2). For a fair benchmark, all tools were executed sequentially using a single CPU core: PADLOC and DefenseFinder processed each genome individually in succession, while PANORAMA first loaded the entire pangenome before sequentially predicting defense systems. Despite this serial execution, PANORAMA significantly outperforms the other tools in runtime, completing analyzes in minutes (S1 Fig & S1 Tab) compared to 24 and 71 hours for DefenseFinder and PADLOC, respectively. This efficiency is achieved by analyzing gene families rather than individual genes, which reduces computational overhead. Additionally, PANORAMA utilizes pyHMMER [27] for HMM alignments, optimizing the workflow by minimizing I/O operations and enabling on-the-fly result filtering. In contrast, other tools use the HMMER software directly [23], which requires post-processing steps.

**Table 2.** Execution performance on *P. aeruginosa* pangenome. Performance comparison of defense system prediction tools on the *P. aeruginosa* pangenome made of 941 genomes, grouped by reference database. All tools were run sequentially on a single CPU with dedicated machine resources. PANORAMA timing excludes pangenome construction

| Tool (version) | Database (version) | #Systems predicted | Run Time (hh:min) | Peak Memory (GB) |
| --- | --- | --- | --- | --- |
| DefenseFinder (1.2.2) | Defense-finder-models (1.2.4) & CasFinder (3.1.0) | 868 | 24:45:22 | 6.84 |
| PANORAMA (1.0.0) |  | 949 | 00:20:05 | 11.22 |
| PADLOC (2.0.0) | PADLOC-DB (2.0.0) | 1038 | 71:27:43 | 3.19 |
| PANORAMA (1.0.0) |  | 1064 | 00:33:33 | 10.77 |

### *Pseudomonas aeruginosa* defense system analysis

#### System prediction and analysis

An exhaustive analysis was conducted on PANORAMA’s defense system predictions, utilizing the DefenseFinder models and the same set of *P. aeruginosa* genomes as for the benchmark. PANORAMA identified a total of 949 systems in the pangenome categorized into 150 distinct models, with restriction-modification (RM) systems being the most abundant (Figure 3). RM systems are present in 84% of genomes, with nearly 300 detected, of which type I systems are the most common, occurring 130 times. The Gabija, CRISPR-Cas, and CBASS system categories follow as the next most prevalent defense systems in genomes, each with a presence rate above 40%. These observations corroborate the study of Johnson *et al.* [28]. At the pangenome level, some system categories are highly prevalent across genomes but are represented by only a few distinct systems. For example, CRISPR-Cas systems appear in 48% of genomes, yet only 13 distinct systems are identified in the pangenome. This is further highlighted by a Shannon entropy calculation, which measures the compositional diversity of system categories (Figure 3C). For CRISPR-Cas systems, the entropy is 1.35, indicating a high degree of conservation in the gene family composition across genomes. Among the most prevalent system categories, others show notable diversity, including RM, Gabija, and RloC. The RloC systems, for example, consist of 22 distinct systems spread across 35% of genomes, with a Shannon entropy of 21.58, indicating considerable variability in their family composition. These predictions are consistent with recent studies and highlight the remarkable diversity of anti-phage immune systems in prokaryotes [11].

**Fig 3.**
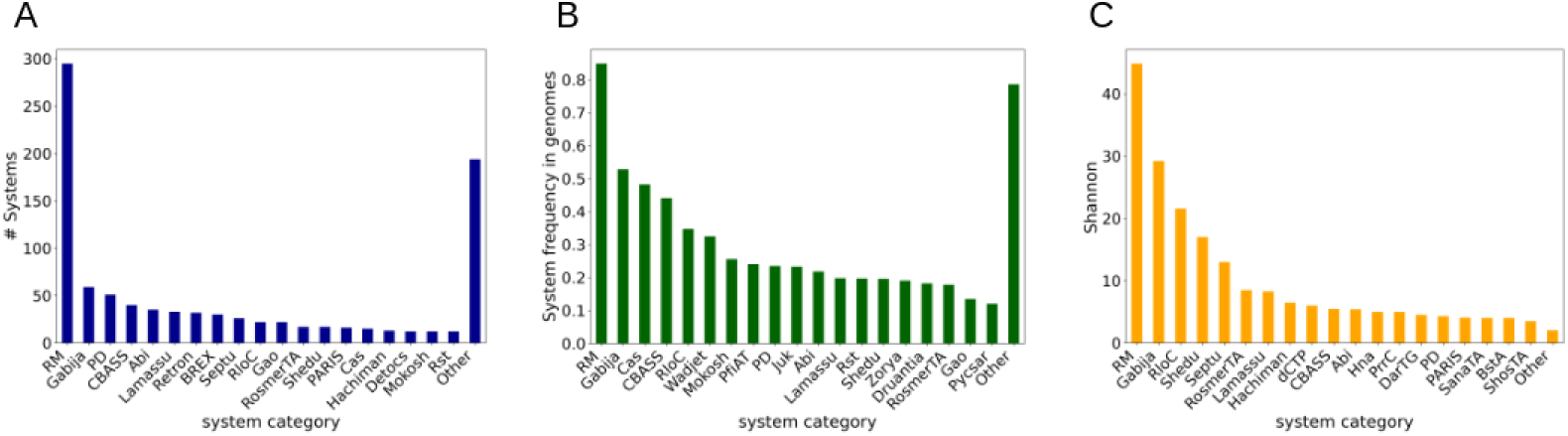
System prediction metrics in *P. aeruginosa*. Systems are grouped by categories on the x-axis and ordered by decreasing values. Only the 19 highest-value system categories are displayed, with others grouped under the ”Other” category. (A) Number of systems found for each category in the pangenome. (B) Relative frequency of system categories in genomes. (C) Shannon entropy of system categories.

#### Defense islands and spots of insertion

PANORAMA systems can be analyzed in conjunction with additional information extracted from the PPanGGOLiN [6] pangenome graph, particularly concerning their association with the variable genome and their localization within RGPs and spots of insertion predicted by panRGP [7]. This enables the identification of defense islands (*i.e.*, variable regions enriched with defense systems) and their hotspots (*i.e.*, frequently occurring insertion sites of defense islands in genomes).

Most defense systems are predicted within the variable (*shell* or *cloud*) genome of *P. aeruginosa* and are located in spots. PANORAMA identified 395 spots containing at least one defense system, representing 40% of all pangenome spots. Among them, 4 spots (7, 6, 45, 69) have a high frequency (*>*25%) and exhibit the highest number of defense systems predicted at the pangenome level (*>*60 systems) (Figure 4). Notably, spots 7 and 6 are the most diverse, harboring 238 and 162 associated systems, respectively (Figure 5). They are mostly composed of RM systems (51% and 56%) but also exhibit a broad diversity of other categories, including BREX (6%), Gabija (5% and 4%), PD (4% in spot 7) and CBASS (4% in spot 6). These two spots were previously identified using a non-automated approach in the study by Johnson *et al.* [28] as core defense hotspots in *P. aeruginosa*, where they were designated CDHS-1 and CDHS-2. This further highlights the value and reliability of PANORAMA in automatically detecting defense islands and their spots. Using PANORAMA, we also identified two additional defense hotspots (spots 45 and 69). Spot 45 contains 110 systems and stands out as the most balanced in terms of system categories; it is also the only hotspot with a notable presence of PARIS systems (8%). Spot 69 contains 75 systems and is dominated by RM systems (like spots 7 and 6), with PrrC systems specifically represented at 7% Although less frequently observed across genomes (*<*20%), spots 61 and 1 display highly diversified system content, comprising 72 and 65 systems, respectively. Both are also rich in RM systems, with Mokosh particularly represented in spot 1 and CBASS and PD systems notably present in spot 61 (≃8%). Finally, spots 4 and 9 are relatively frequent across genomes (≃30%) but contain few distinct systems, 31 and 10, respectively. These results highlight the potential of PANORAMA to provide a comprehensive landscape of defense systems in a species, enabling pangenome-scale analysis and the identification of defense islands along with their hotspots of insertion.

**Fig 4.**
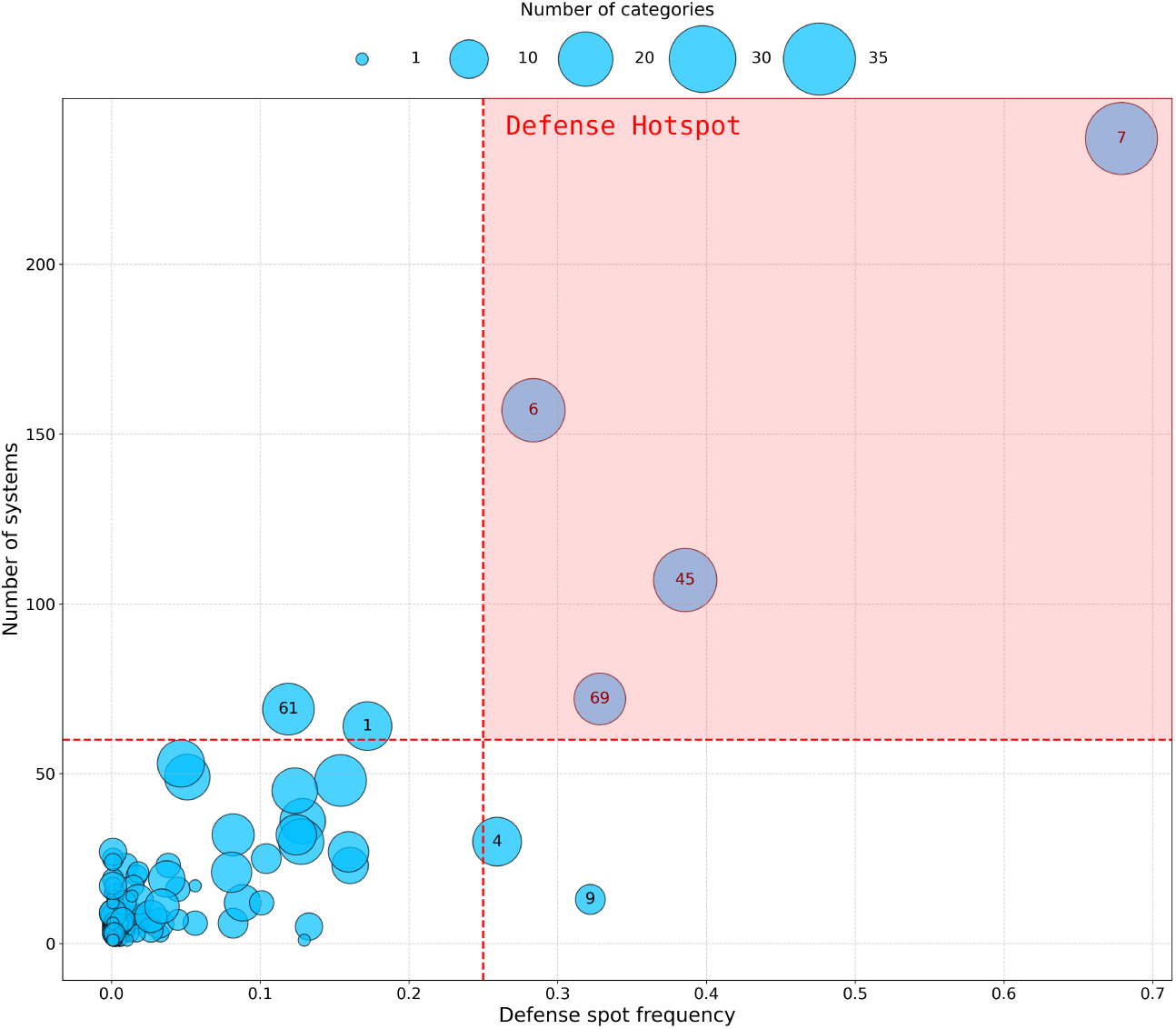
System diversity and defense spot frequency in *P. aeruginosa*. This bubble plot displays the distribution of defense spots identified by PANORAMA, based on their frequency in genomes (x-axis) and the total number of defense systems identified within each spot at the pangenome level (y-axis). The size of the bubbles is proportional to the number of distinct system categories represented in each spot (legend at top shows scale), and the numbers within bubbles indicate spot identifiers.

**Fig 5.**
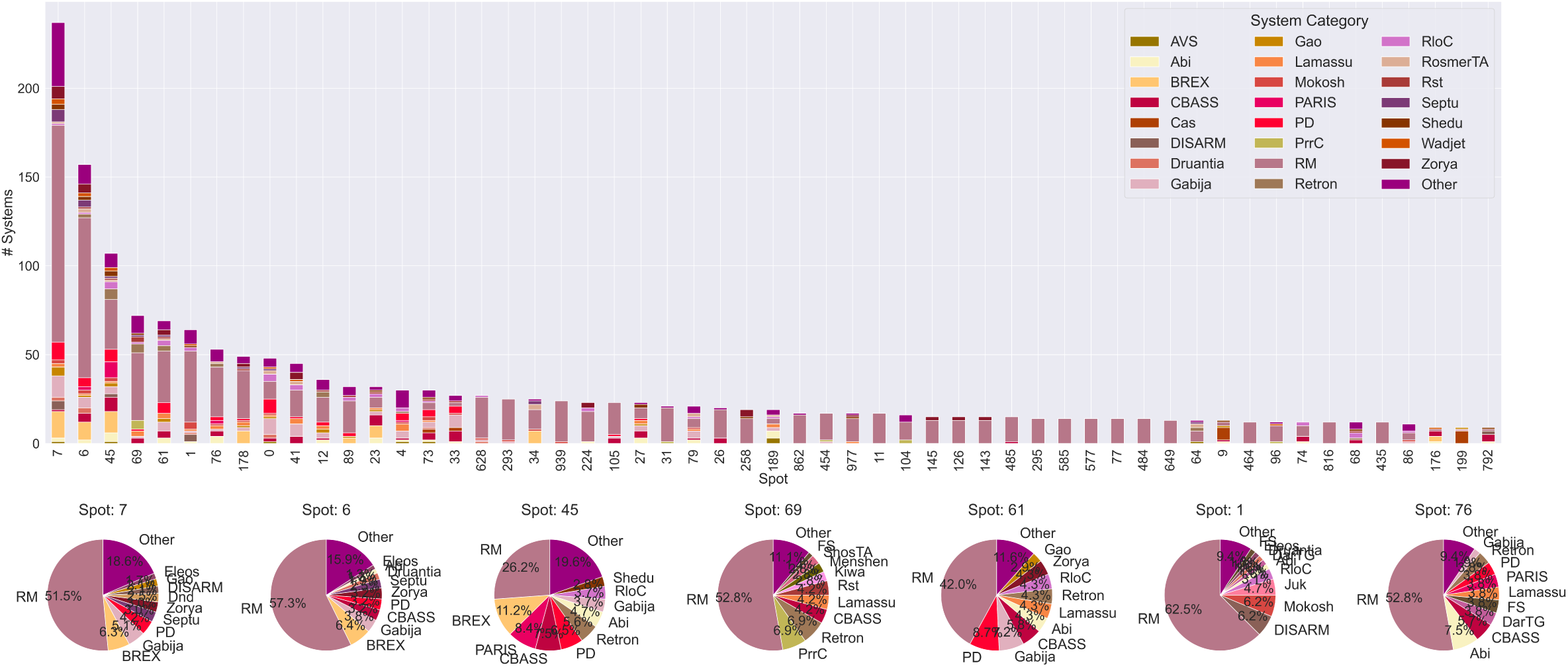
Defense systems within insertion spots of the *P. aeruginosa* pangenome. The bar plot (top) displays the number of predicted defense systems in the *P. aeruginosa* pangenome for each spot. Only spots with at least 10 systems are displayed. The pie charts (bottom) illustrate the system category composition for the six major spots.

### Pangenome comparison of Enterobacteriaceae defense arsenal

To demonstrate the comparative functionalities of PANORAMA (*i.e.*, PanCompare workflow), the pangenomes of four Enterobacteriaceae species were analyzed for defense system and spot prediction. This dataset represents more than 6,000 genomes from *Citrobacter freundii*, *Salmonella enterica*, *Klebsiella pneumoniae*, and *Escherichia coli* species (Table 3). The distribution of systems between species and their association with conserved spots were studied. Defense systems were predicted using DefenseFinder models.

**Table 3.** Pangenome statistics across bacterial species. Summary statistics for five bacterial species analyzed in this study.

| Species | Genomes | Genes | Families | Edges | Persistent | Shell | Cloud | RGPs | Spots |
| --- | --- | --- | --- | --- | --- | --- | --- | --- | --- |
| *C. freundii* | 79 | 385611 | 16865 | 24936 | 3780 | 2954 | 10131 | 3863 | 254 |
| *E. coli* | 3083 | 14447407 | 44278 | 140743 | 3018 | 7174 | 34086 | 240163 | 2036 |
| *K. pneumoniae* | 1659 | 8851686 | 32685 | 75073 | 4155 | 5085 | 23445 | 67678 | 778 |
| *P. aeruginosa* | 941 | 5811655 | 34058 | 61841 | 4925 | 6768 | 22365 | 44935 | 984 |
| *S. enterica* | 1380 | 6222501 | 23941 | 46535 | 3474 | 4080 | 16387 | 65920 | 458 |

#### Defense system distribution in the four species

A total of 344, 452, 988, and 1470 defense systems were predicted from the pangenomes of *C. freundii*, *S. enterica*, *K. pneumoniae*, and *E. coli*, respectively. In addition to textual outputs, PANORAMA automatically generates a heatmap that displays system occurrences across the compared pangenomes (see S2 Fig). Among the different categories, RM systems are the most abundant in the four species, accounting for 30% to 45% of the systems found. Following RM systems, PD systems are the next most prevalent, representing 5% to 7% of the systems within Enterobacteriaceae (Figure 6). Other notable system categories, such as Retron, CBASS, Gao, BREX, and Abi, are also relatively abundant across all pangenomes. Next, we evaluated the species-specificity of each system category by computing enrichment factors (S3 Fig). Our findings indicate that certain categories, Azaca and Dnd in *S. enterica*, Juk in *K. pneumoniae*, and Bunzi and RADAR in *C. freundii*, exhibit enrichment factors above 3, highlighting their preferential association with these species. Tiamat systems are only found in *S. enterica* and *C. freundii*. Interestingly, *E. coli* does not show any systems in higher abundance compared to other species. Beyond the large number of genomes used to construct the pangenome, this pattern may reflect that it includes strains exposed to a wider diversity of phages, a consequence of its commensal lifestyle in the gut.

**Fig 6.**
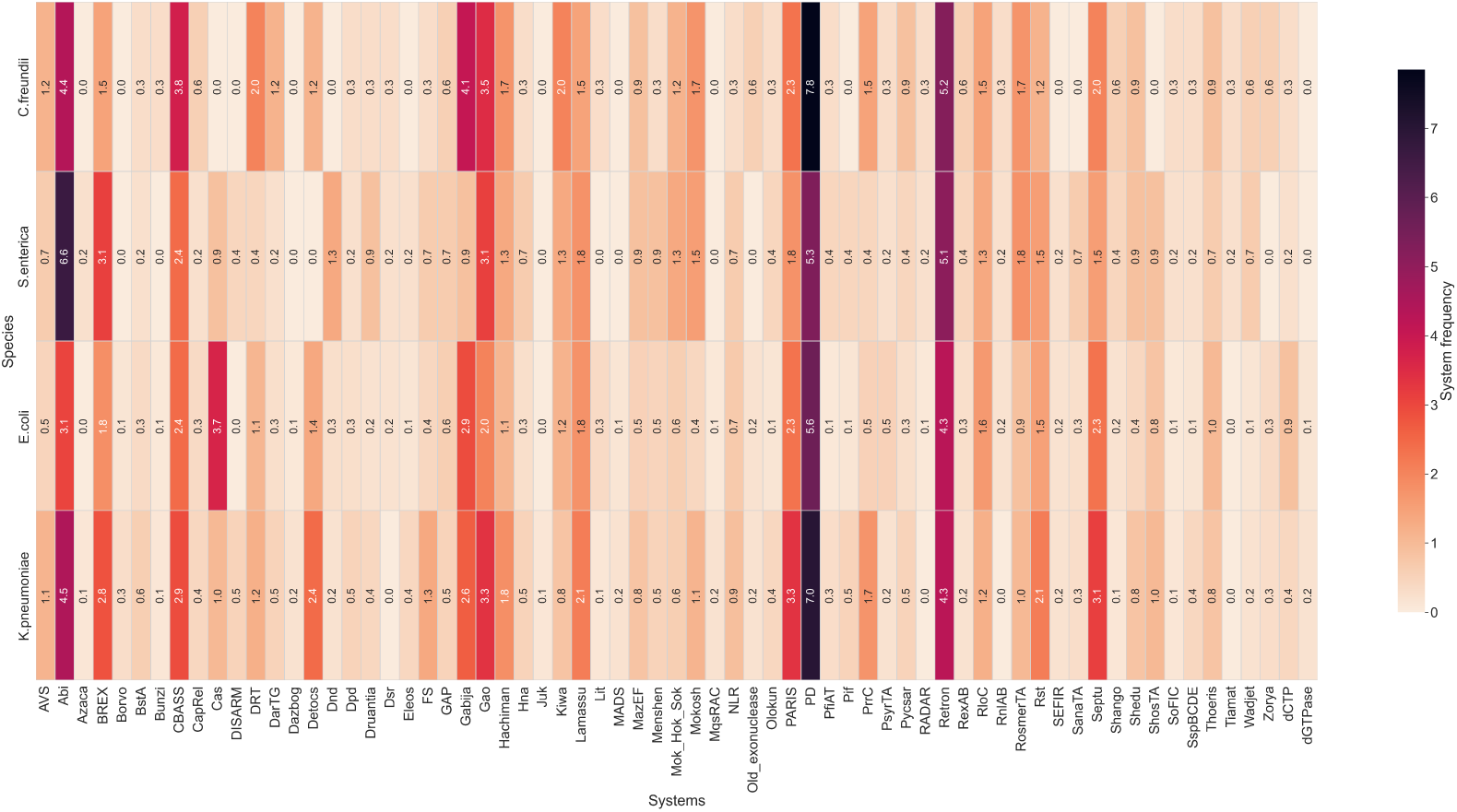
Relative frequency of system categories in Enterobacteriaceae pangenomes. The relative frequencies are expressed as a percentage. The RM system category was omitted to facilitate interpretation.

#### Identification of conserved spots

We identified 150, 291, 635, and 1664 spots with at least one defense system in *C. freundii*, *S. enterica*, *K. pneumoniae*, and *E. coli*, respectively. The most defense system-rich spot in our analysis is spot 35 in *E. coli* (S4 Fig), which contains more than 450 systems and corresponds to the tRNA-LeuX genomic island. This island, previously characterized for harboring multidrug resistance genes [29], is revealed here to also serve as a major phage defense hotspot, demonstrating the dual protective role of this genomic region.

With PANORAMA, we searched for similar spots based on their related gene families, using a gene family repertoire relatedness (GFRR, see Pangenome comparison workflow) threshold of at least 60%. This threshold guarantees at least one similar gene family on each side of the border. As shown in Figure 7, we identified 102 clusters of similar spots, encompassing 232 spots that each share at least one neighboring gene family composition with another spot across the Enterobacteriaceae pangenomes.

**Fig 7.**
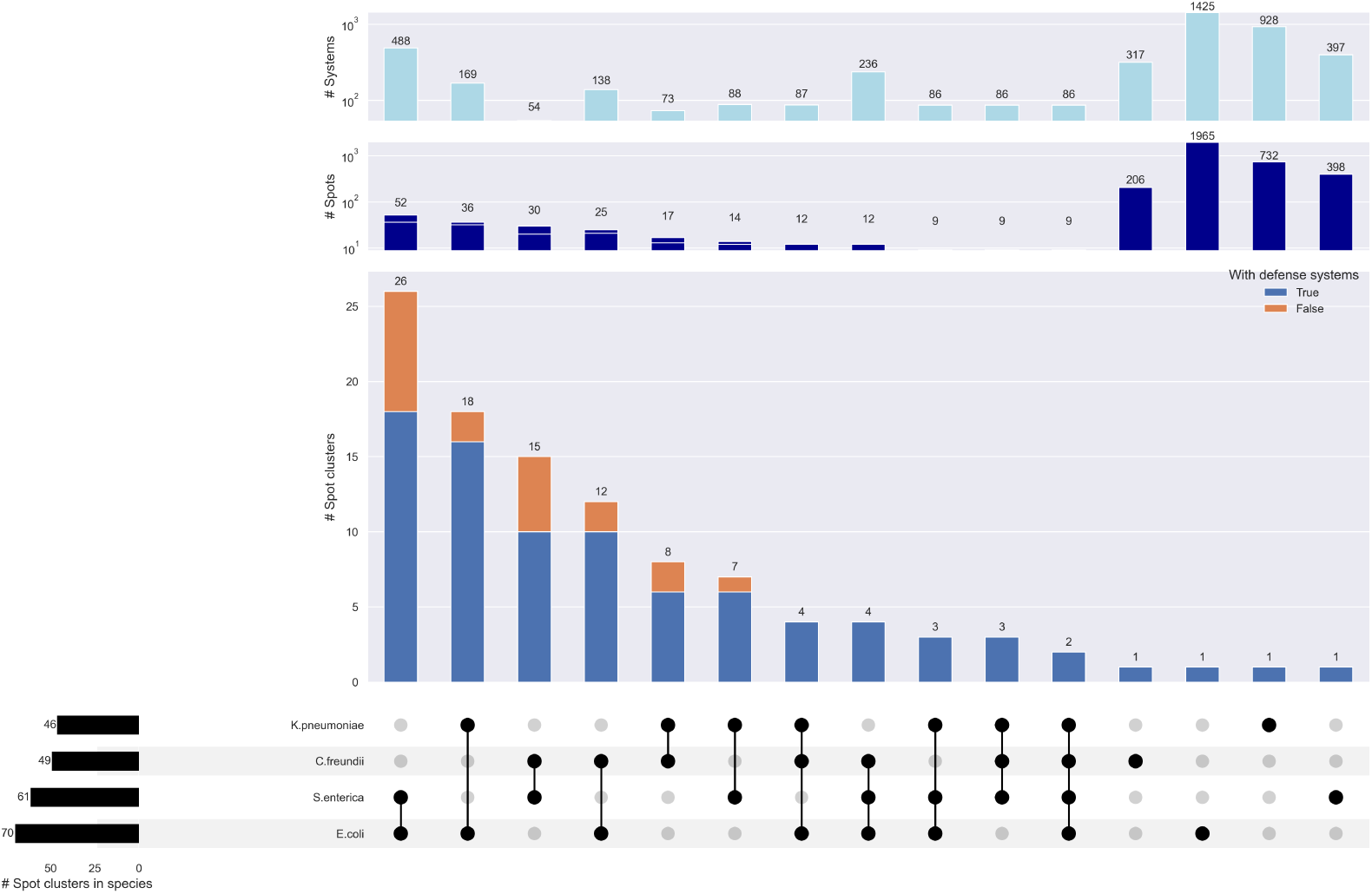
Sharing of spot clusters across four Enterobactariaceae species and their association with defense systems. The UpSet plot shows the number of spot clusters shared across the four compared species, with stacked bars to indicate whether they contain defense systems (in blue) or not (in orange). The two top bar plots represent spot and defense system abundance metrics within spot clusters on a logarithmic scale.

*E. coli* is the species that shares the most spots with others (71), followed by *S. enterica* (60), then *C. freundii* (48), and *K. pneumoniae* (46). Proportionally to its number of spots, *C. freundii* is the species with the most similar spots, with 18% of its spots similar to the other pangenomes. The 2 species with the most common spots are *E. coli* and *S. enterica*, with 67 common spots. PANORAMA can then be used to highlight spots of insertion at a higher taxonomic rank than the species. These could be areas of interest for research into shared evolution or exchanges between these species.

#### Defense spot identification and conservation

Using the comparative functionalities of PANORAMA, we identified 99 clusters of similar spots conserved in at least two of the four Enterobacteriaceae species (Figure 7). As might be expected, given their phylogenetic proximity, *E. coli* and *S. enterica* share the most common spots (*i.e.*, 34 spot clusters, 26 of which are found only in these two species). About half of the spot clusters (n=47) are associated with a defense system in at least one species, comprising a total of 520 distinct defense systems among the 3,265 identified in the four species pangenomes. Of these, there is only one spot cluster (cluster 58) conserved in all pangenomes, containing 34 defense systems.

*E. coli* and *S. enterica* harbor the highest number of defense systems (263 systems) across their specific spot clusters (7 clusters). One of these clusters (spot cluster 3062) includes spot 86 in *S. enterica* and spot 175 in *E. coli*. These spots rank, respectively, eighth and sixth in terms of the number of defense systems (S4 Fig), with 32 in *S. enterica* and 216 in *E. coli*, and can therefore be considered as potential defense islands or even defense hotspots. Analyzing their composition reveals notable similarities (S5 Fig). Both spots are mainly composed of RM systems (55% in *S. enterica*, 85% in *E. coli*), followed by BREX, CBASS, and PrrC, which are generally not phage-specific. These findings support the hypothesis that defense systems may have been exchanged between the two species from this conserved hotspot. Interestingly, spot 175 in *E. coli* corresponds to a previously identified defense hotspot (spot 4) by Hochhauser *et al.* [22]. The expanded system diversity and abundance we observe, compared to the previous study, reflect our larger genome dataset.

Examining the system category diversity of other spot clusters (Figure 8), many clusters are predominantly composed of RM systems. Some clusters are dedicated to a single system category, such as BstA in cluster 2320, while others, like the previously mentioned cluster 58, are more diverse, bringing together systems from all four studied species.

**Fig 8.**
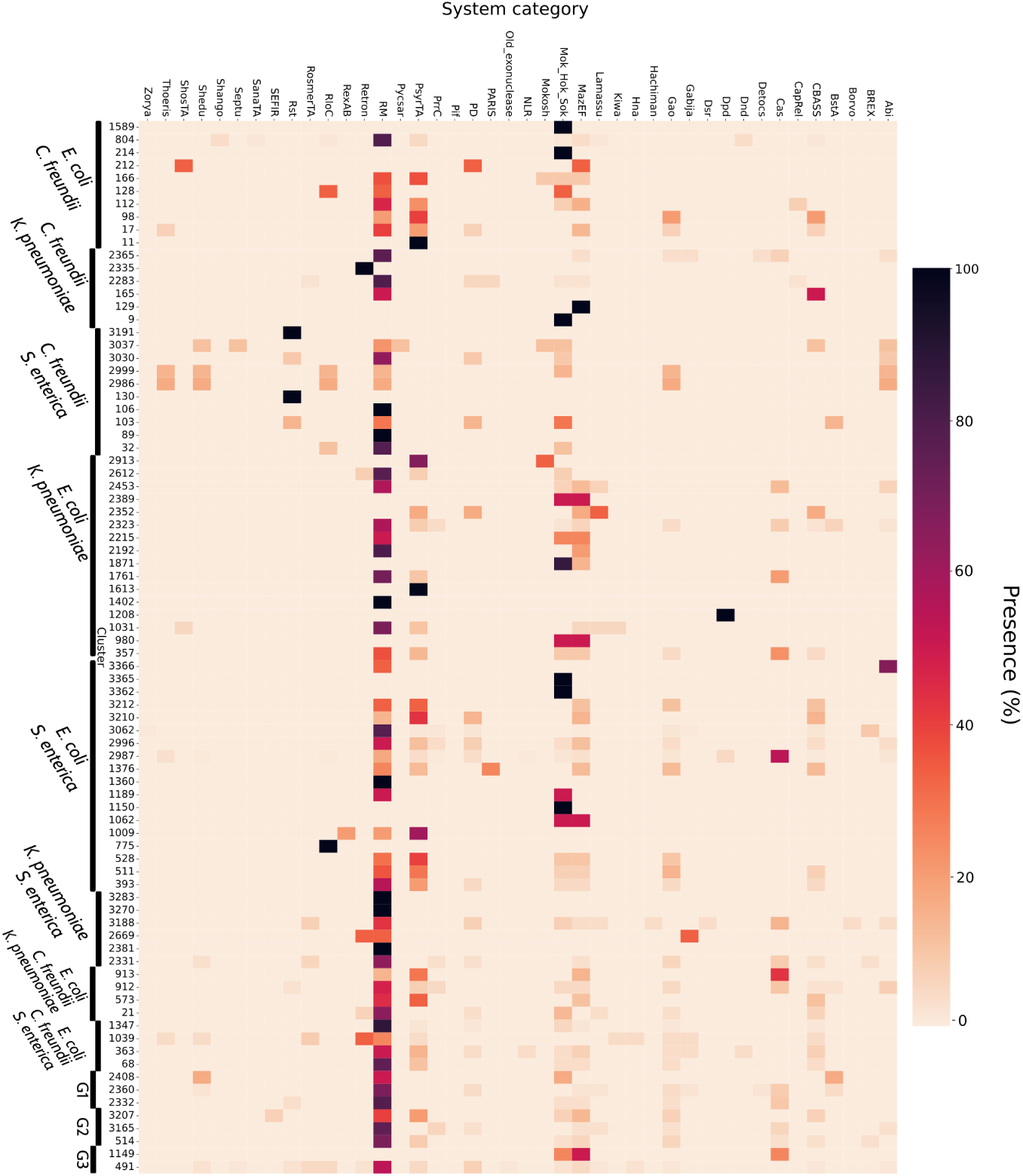
Relative frequency of system categories in each spot cluster. The heatmap presents the relative frequency (%) of system categories within each spot cluster. Species associated with each spot cluster are indicated next to the corresponding cluster identifiers. G1: *C. freundii*, *K. pneumoniae*, *S. enterica*; G2: *E. coli*, *K. pneumoniae*, *S. enterica*; G3: *C. freundii*, *E. coli*, *K. pneumoniae*, *S. enterica*

Beyond their illustrative purpose, the analyzes presented here highlight the ability of PANORAMA’s comparative functionalities to identify conserved defense islands across species, providing valuable insights into the evolution of defense systems and their mechanisms of acquisition.

## Conclusion

In this work, we introduced PANORAMA, a novel open-source framework for the prediction and analysis of macromolecular systems across prokaryotic pangenomes. Unlike existing tools that operate on individual genomes, PANORAMA leverages the pangenome graph structure provided by PPanGGOLiN to perform system-level analyzes at both species and interspecies scales. It utilizes rule-based models inspired by those in MacSyFinder, but adapted to pangenomes, enabling the flexible and accurate identification of diverse functional systems, such as defense mechanisms.

Benchmarking against established tools, such as DefenseFinder and PADLOC, demonstrated that PANORAMA achieves comparable prediction accuracy while reducing computational demands, making it particularly well-suited for large-scale analyzes involving hundreds or thousands of genomes.

Beyond its performance, PANORAMA introduces key methodological innovations. Notably, its graph-based approach enables the detection of conserved genomic contexts by identifying gene families that consistently co-occur across multiple genomes. This strategy allows for the robust identification of evolutionarily maintained system components, even when their genomic proximity is disrupted in some genomes by rearrangements, insertions, or assembly fragmentation. Consequently, PANORAMA offers greater flexibility than traditional genome-based tools in identifying atypical genomic architectures.

Applied to hundreds of genomes of a species, PANORAMA can quickly identify the different systems and their occurrences in individual genomes. A notable strength lies in its ability to associate systems with regions of genomic plasticity and hotspots of insertion, allowing for the identification of genomic islands enriched with systems and their preferred integration sites. In *P. aeruginosa*, four major hotspots were identified, including two that correspond to previously published core defense hotspots.

To further illustrate the comparative functionality of PANORAMA, we analyzed the defense repertoires of over 6,000 genomes from four Enterobacteriaceae species. PANORAMA revealed both conserved and species-specific system categories and was able to cluster insertion spots based on shared gene family content. This enabled the identification of conserved defense islands across phylogenetically related species, offering insights into the evolutionary conservation of these systems.

Altogether, PANORAMA provides a robust and extensible framework for system detection and comparative analysis at the pangenome level. An important advantage of PANORAMA is that its predictions are inherently produced at the pangenome scale: whereas genome-based tools require the aggregation and harmonization of hundreds or thousands of individual genome-level outputs to obtain species-wide summaries, PANORAMA directly provides unified results through its graph-based representation. Its flexible modeling format, compatibility with existing system model databases, and integrative workflows make it a valuable tool for microbiologists, bioinformaticians, and evolutionary biologists interested in understanding the distribution, function, and evolution of macromolecular systems. As the number of available genomes continues to grow, tools like PANORAMA will become increasingly critical in unveiling the complex architectures of microbial defense and other macromolecular systems.

Looking ahead, we plan to expand PANORAMA’s detection capabilities by incorporating anti-defense system models [30] from DefenseFinder (v1.3.0), enabling analysis of the co-evolutionary dynamics between defense and anti-defense mechanisms. We also plan to incorporate additional rule-based approaches to predict metabolic modules, detecting genomic contexts encoding enzymes involved in the same pathway using comprehensive databases such as KEGG [31] or MetaCyc [32], as well as develop specific rules for detecting clusters of Carbohydrate-Active Enzymes (CAZymes) involved in polysaccharide biosynthesis and degradation. All these model collections will be made publicly available on GitHub (https://github.com/PANORAMA-models) and, following the approach of other tools in the field, we will implement functionality within PANORAMA to automatically download the latest version of these models, simplifying the functional analysis of pangenomes. Furthermore, the pangenome graph framework provided by PANORAMA offers a promising avenue to explore genomic context and uncover novel systems. By incorporating fuzzy functional predictors based on structural similarity or protein language models, we hope to detect new systems composed of functionally analogous components co-localized within the pangenome graph.

## Materials and methods

### Overview of PANORAMA software

PANORAMA is an open-source Python3 software (version ≥3.10 required) that enables pangenome-scale detection and analysis of prokaryotic macromolecular systems and pangenome comparison to identify similar structures based on gene family content. The tool is freely available on GitHub (https://github.com/labgem/PANORAMA) and is distributed under the CeCILL 2.1 license. Built on the PPanGGOLiN API version 2.3.0, PANORAMA inherits its core concepts and definitions for pangenome partitions, gene families, RGPs, and spots of insertion [6, 7]. The software can be easily installed via bioconda [33] (https://anaconda.org/bioconda/panorama), ensuring reproducible deployments across different computing environments. Comprehensive documentation is available online at https://panorama.readthedocs.io/latest/, including detailed analysis workflows, parameter descriptions, usage examples, and a step-by-step guide for creating custom detection models.

### Pangenome system detection workflow

To predict macromolecular systems, PANORAMA employs rule-based models that analyze the presence/absence of specific functions predicted from pangenome gene families while considering constraints on their genomic organization. The PanSystem workflow begins with gene family annotation using HMM libraries, followed by the identification and evaluation of genomic contexts within the pangenome graph based on system model rules (Figure 9). Finally, the predicted systems at the pangenome level are mapped onto individual genomes to assess their presence and gene content.

**Fig 9.**
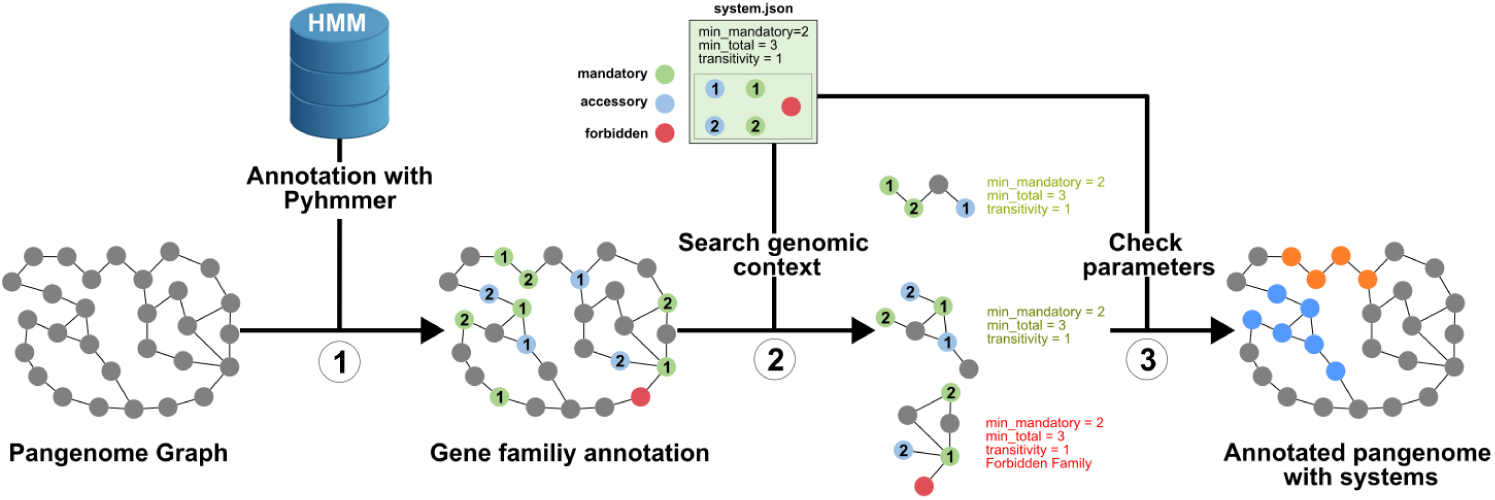
PANORAMA PanSystem workflow. The detection workflow for macromolecular systems consists of multiple sequential steps: (1) gene family annotation using HMM profiles, (2) detection of genomic contexts from the pangenome graph containing annotated families, (3) checking model rules, (4) projection of systems on genomes.

#### System modeling

Models used by PANORAMA for predicting macromolecular systems are similar to those of MacSyFinder [9] but differ in some aspects (Figure 10). The primary components of a system *Model* are *Families* (*i.e.*, isofunctional protein families) instead of genes, and an additional hierarchical level is introduced to represent *Functional Units*. A *Functional Unit* is defined as a set of *Families* that work together to perform a necessary function for the system. Several *Functional Units* may be required for a system to operate effectively. This new level provides a more detailed and accurate description of systems, allowing the use of distinct rules for presence/absence and genomic organization, both for the *Families* of a *Functional Unit* and between *Functional Units*. In PANORAMA, we also introduce the concept of canonical *Models*, which represent a more relaxed definition of systems. In PADLOC [10], these models are identified by the keyword ”other” in their names. While these systems may not have been experimentally validated and might not be functional, their predictions can be valuable for identifying new systems or potential variants. During system detection, priority is given to non-canonical models. A canonical model is predicted only if its *Families* are not already associated with a non-canonical model.

**Fig 10.**
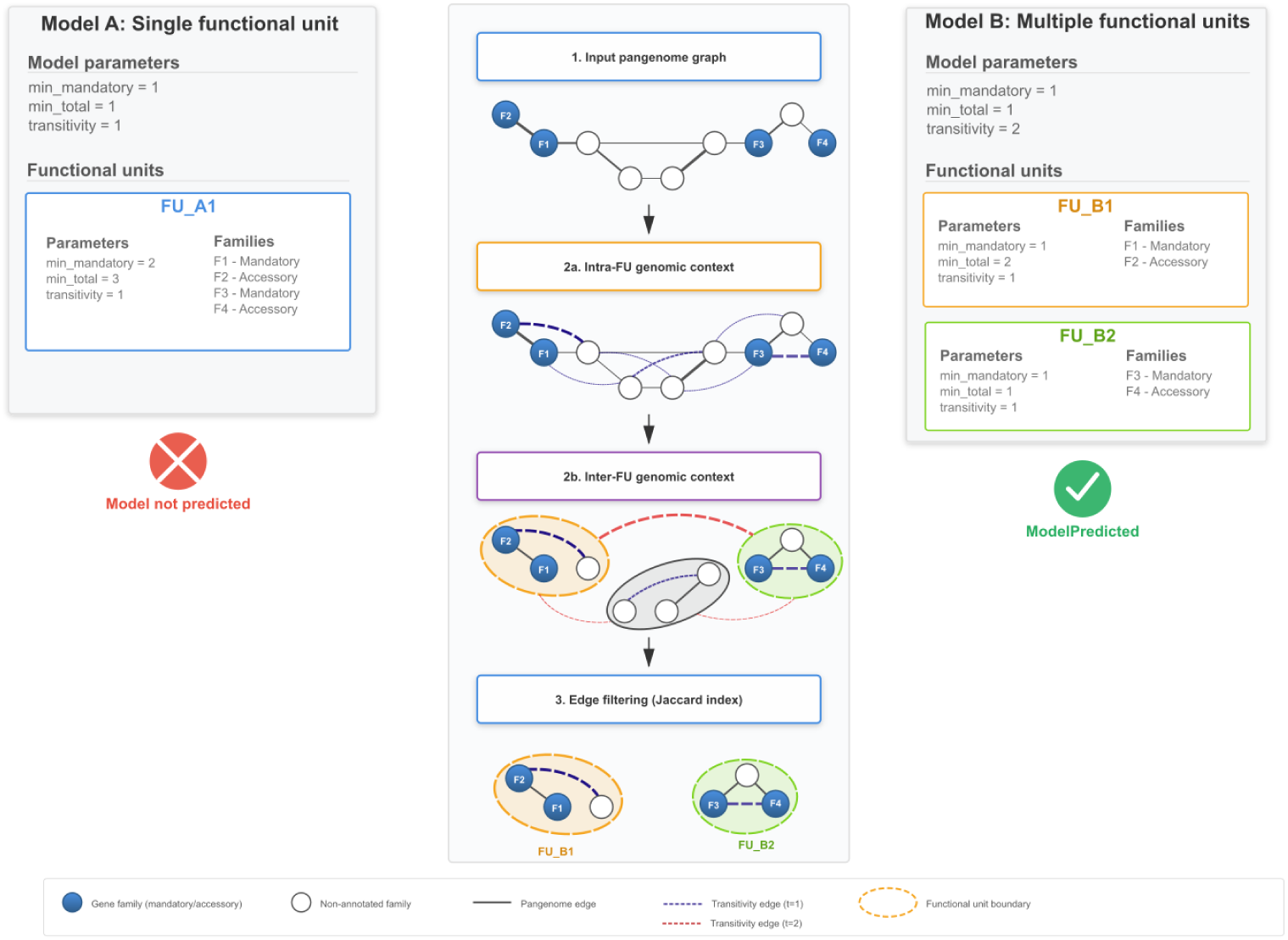
PANORAMA system modeling. Two modeling approaches are compared: Model A (left) uses a single functional unit with parameters that fail to detect the structure (*min mandatory* = 1, *min total* = 1, *transitivity* = 1), while Model B (right) employs multiple functional units with increased transitivity (*transitivity* = 2), successfully identifying FU B1 (orange) and FU B2 (green). Blue nodes represent gene families within the model (mandatory or accessory), white nodes indicate gene family without function for the system, solid black lines show pangenome edges, and dashed lines represent transitivity edges connecting genes at specified graph distances. Colored dashed ovals delineate predicted functional unit boundaries.

For presence/absence constraints, each *Functional Unit* and *Family* is categorized as *mandatory*, *accessory*, *neutral*, or *forbidden*. *Mandatory* elements are essential for the system. *Accessory* components are dispensable. They contribute to the system, but may not be identified in all system variants due to rapid evolution or the absence of homology. *Forbidden* elements are incompatible with system functionality. They can help differentiate systems with shared components or distinguish inhibited systems. Although *Neutral* components are considered associated functions, they are not used to assess system predictions. However, they can serve as intermediaries in genomic context analysis by linking mandatory or accessory elements. To predict a system, quorum rules on component presence/absence are defined by two parameters: *minimum mandatory* (the minimum number of mandatory elements required) and *minimum total* (the minimum number of both mandatory and accessory elements). These parameters are specified at two levels: at the *Model* level, to assess the presence of *Functional Units*, and at the *Functional Unit* level, to evaluate predicted *Families*. Constraints on the genomic organization of a system are governed by a *transitivity* parameter, which defines the maximum genomic distance between gene families in the pangenome graph corresponding to the system components (see Pangenome context extraction).

#### Functional annotation of pangenome gene families

To annotate the pangenome graph, PANORAMA utilizes HMM libraries, in which each HMM represents a specific *Family* defined in the system model. Protein sequences of the family’s representative genes are aligned to the HMM profiles using the hmmsearch method from the pyHMMER Python library [27]. PANORAMA also offers the ability to use the consensus protein sequence of gene families. It is determined by performing a multiple sequence alignment (MSA), excluding fragmented genes, and then computing the consensus sequence with pyHMMER. To validate the alignments, a metadata file can be linked to an HMM library, specifying thresholds for each HMM based on various criteria, such as alignment coverage on the sequence or profile, score, e-value, and independent e-value. Alternatively, a global threshold can be applied to all HMMs. An option is also available to keep only the *n* best hits. Gene family annotation can also be performed externally using alternative prediction methods. In this case, a TSV file can be provided to describe the functions associated with each gene family.

#### Pangenome context extraction

For each system, connected components between gene families corresponding to model families (hereafter called target families) are searched for in the pangenome graph. According to the model rules on genomic colocalization of system components (*i.e.*, the *transitivity* parameter), additional edges are added to the pangenome graph between families separated by fewer than *t* genes in the corresponding genomes. This corresponds to a partial transitive closure on the pangenome graph, enabling the connection of two target families even if their genes are not directly adjacent in the genomes. Next, a Jaccard index-based criterion (Equation 1) is computed on edges to evaluate genomic context conservation (*i.e.*, synteny conservation). For two families *a* and *b* connected by an edge *e_a,b_*, the index *J_a,b_* is defined as the ratio of the number of genomes in which the edge *e_a,b_* exists to the number of genomes in which at least one of the two gene families is present (|gen*_a_* ∪ gen*_b_*|). To enhance the detection of systems present in a limited number of genomes, this index is computed locally, considering only the genomes that contain the target families of the connected component rather than all the genomes of the pangenome. Edges having a Jaccard index below a defined threshold (*J_a,b_ <* 0.8 by default) are removed. The connected components with their remaining nodes are then evaluated in terms of presence/absence rules to validate system predictions. For each connected component, this process, which combines edge filtering and presence/absence rule validation, is applied iteratively, starting with the largest set of target families observed in a genome and then exploring other sets of target families, from largest to smallest, if they are not already included in a predicted system.

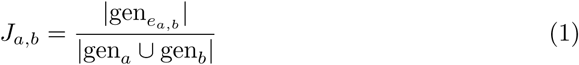

Finally, the connected components corresponding to predicted systems at the pangenome level are mapped back onto individual genomes. Their occurrence in individual strains is assessed based on presence/absence and colocalization rules. They are classified into 3 distinct categories: *strict*, *extended*, or *split*. A system is flagged as *strict* if the *transitivity* parameter is respected among the genes belonging to the system. If additional genes of the connected component are found among those encoding the system, it is classified as *extended*. Indeed, applying transitive closure to the pangenome graph enables the detection of additional genes conserved with those of the system. This provides a more relaxed definition of colocalization rules than the strict intergenic space constraints employed by MacSyFinder or PADLOC. Lastly, *split* systems are those in which system genes are separated by non-member genes of the connected component detected at the pangenome level or are located on different contigs. Such fragmentation can occur in a subset of genomes due to rearrangements, insertion events (*e.g.*, insertion sequences), or assembly breaks.

### Pangenome comparison workflow

The PanCompare workflow of PANORAMA enables the comparison of pangenomes and the identification of similar elements across species, such as macromolecular systems or spots of insertion. These elements are represented as sets of gene families. The first step is to cluster together all the gene families from the different pangenomes in order to identify groups of homologous gene families (GHGFs). Then, a Gene Family Repertoire Relatedness score (GFRR) for each pair of elements is computed. Finally, a community clustering algorithm is applied to identify clusters of elements sharing similar gene family content.

#### Gene families clustering

The process involves the extraction of representative protein sequences of gene families from all pangenomes. These sequences are then clustered using the MMSeqs2 cluster command [34] to obtain GHGFs sharing at least 50% of amino acid identity (*–min-seq-id* parameter) with an alignment coverage of 80% (*-c* parameter) by default. The following additional parameters are used: *–max-seqs* 400 *–min-ungapped-score* 1 *–kmer-per-seq* 80 *–alignment-mode* 2 *–cluster-mode* 1.

#### Gene Family Repertoire Relatedness score

To compare the family content of two pangenome elements, two GFRR scores are computed using GHGFs. For two pangenome elements, *a* and *b*, with their GHGFs content denoted as *GHGF_a_*and *GHGF_b_*, the minimal GFRR score, *minGFRR_a,b_*, is defined as the ratio of the number of gene families belonging to the same GHGFs in *a* and *b* to the minimum number of gene families between *a* and *b* (Equation 2). Similarly, the maximal GFRR score, *maxGFRR_a,b_*, is computed using the maximum number of gene families between *a* and *b* (Equation 3).

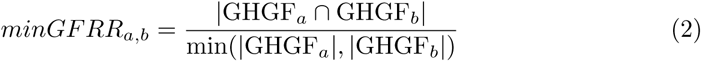

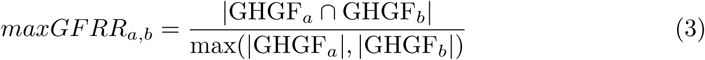

GFRR score computation can be applied to predicted macromolecular systems or spots of insertion. For spots, the two sets of gene families corresponding to spot borders are merged to compute the GFRR scores.

#### Pangenome element clustering

To identify similar elements (*i.e.*, spots of insertion or systems) between pangenomes, GFRR scores are computed for each pair of elements. A graph is then constructed, where nodes represent pangenome elements and edges are weighted by their corresponding GFRR scores. Edges are filtered according to a GFRR threshold (*min_GF RR_ >*= 0.6 by default) to ensure strong similarity between connected elements. Finally, a Louvain algorithm [35], using the NetworkX library [36] implementation with GFRR weights on edges, is applied to the graph to identify non-overlapping communities corresponding to groups of similar elements between pangenomes. This functionality was used to identify conserved spots of insertion between species containing defense systems. A system is associated with a spot if all of its gene families are part of RGPs found within the boundaries of that spot.

### Data, benchmark, and metrics

#### Genomic data and defense system prediction

Complete genomes of five species were downloaded from NCBI RefSeq [37] and analyzed for defense system prediction using DefenseFinder (v1.2.2 with 239 models of v1.2.4 + 43 CasFinder models of v3.1.0), PADLOC (v2.0.0, 385 models of v2.0.0), and PANORAMA (v1.0.0). The dataset includes *Pseudomonas aeruginosa* (941 genomes) and four well-studied Enterobacteriaceae species: *Citrobacter freundii* (79 genomes), *Escherichia coli* (3,083 genomes), *Klebsiella pneumoniae* (1,659 genomes), and *Salmonella enterica* (1,380 genomes). Before running PANORAMA, the pangenomes of the five species were obtained using PPanGGOLiN v2.3.0 with default parameters while keeping the original RefSeq annotations. Pangenome metrics, including the number of families categorized as *persistent*, *shell*, or *cloud* genomes, as well as the number of predicted RGPs and spots of insertion, are summarized in Table 3. These metrics are also available through the info command of PANORAMA. Models from DefenseFinder and PADLOC (n=667), along with their associated HMMs (n=6,272), were converted to PANORAMA format using its internal conversion utility.

#### Benchmark protocol

The *P. aeruginosa* genome dataset was used to evaluate PANORAMA’s defense system predictions against the reference tools, DefenseFinder v1.2.2 and PADLOC v2.0.0, using their respective system models and HMMs. To ensure consistency in comparison, the predictions from DefenseFinder and PADLOC, originally consisting of gene sets associated with defense systems, were mapped to the pangenome gene families by linking each identified gene to its corresponding family. Then, we define a True Positive (TP) as a gene family assigned to the same defense system by both PANORAMA and the reference tool. Gene families associated with a system by the reference tool but not predicted by PANORAMA are classified as False Negatives (FN). Gene families assigned to a different system or not predicted by the reference tool are considered False Positives (FP). To assess the performance of PANORAMA, we computed precision, recall, and F1-score using the following equations:

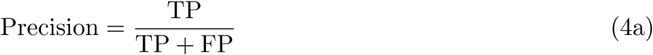

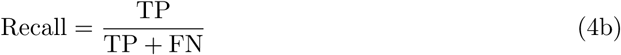

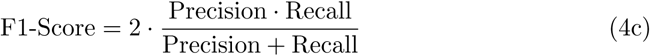

The benchmark was conducted on a dedicated Linux server equipped with two Intel® Xeon® Gold 6150 processors (36 cores), 376 GiB of DDR4 RAM, running CentOS v7.9.2009 with kernel 3.10.0 and Python 3.10. To ensure fair comparison and consistent resource availability, the entire machine was reserved for each tool execution. DefenseFinder and PADLOC were run sequentially, processing one genome at a time on a single CPU core, with exclusive access to all available memory. Similarly, PANORAMA was run with the *–threads* parameter set to 1 with dedicated machine access. To evaluate execution performance, total run time and peak memory usage were measured for each tool. For consistency in comparison, the pangenome construction step performed by PPanGGOLiN was not included in PANORAMA execution metrics, as pangenomes are inputs of PANORAMA, analogous to genomes for PADLOC and DefenseFinder.

#### System category diversity

To assess the compositional diversity of systems within a given category (*e.g.*, Restriction Modification, Gabija, CRISPR-Cas) in a pangenome, Shannon entropy is calculated as follows:

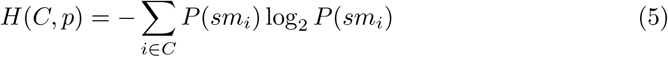

Where:

- *H*(*C, p*): the Shannon entropy of a system category *C* in a pangenome *p*.
- *P* (*sm_i_*): probability of a system model *sm_i_* of *C* occuring in *p*, calculated for each system model as the ratio of the number of gene families of *p* associated with *sm_i_* to the total number of gene families across all systems of *C*.

Higher entropy indicates greater diversification of the gene families comprising the systems within a given category.

#### System category enrichment

To determine the specificity of a system category in a set of pangenomes, an enrichment factor is computed as follows:

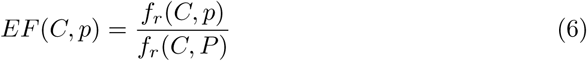

Where:

- *C*: a system category.
- *P*: a collection of pangenomes *p*.
- *f_r_*(*C, p*): the relative frequency of the system category *C* in the pangenome *p*, calculated as the ratio of the number of systems of *C* predicted in *p* to the total number of systems in *p*.
- *f_r_*(*C, P*): the relative frequency of the system category *C* in *P*, calculated as the ratio of the number of systems of *C* predicted in all *p* ∈ *P* to the total number of systems across all *p* ∈ *P*.

An enrichment factor above 1 indicates that a system category appears n times more in a specific pangenome than in the entire collection of pangenomes.

## Supporting information

**S1 Fig.**
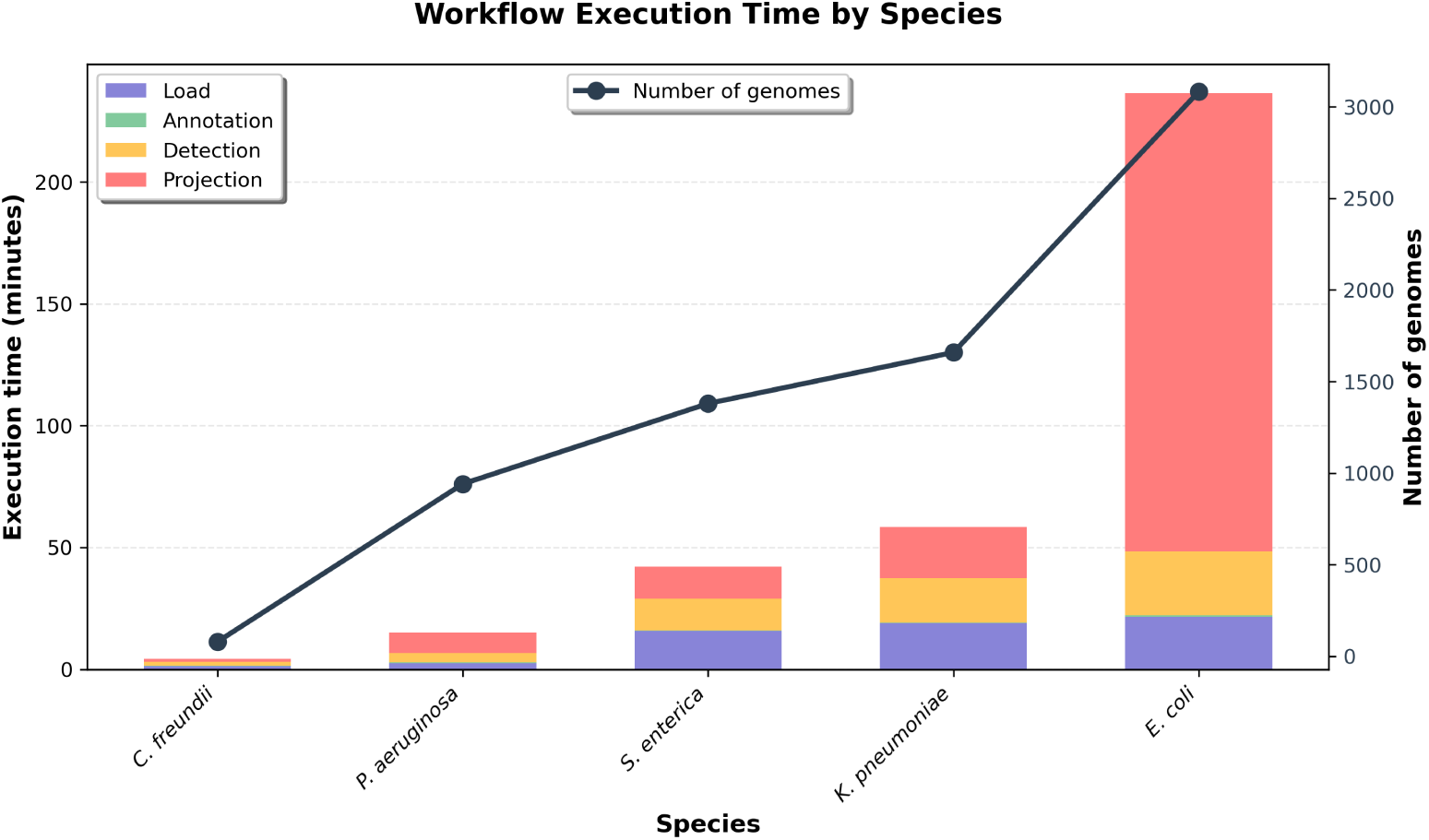
Computational performance of the analysis pipeline across bacterial species. Stacked bar chart showing the execution time (in minutes) for each step of the workflow across species with varying numbers of genomes (black line, right y-axis). The four pipeline steps are: Load (blue) - time to load the pangenome data; Annotation (green) - time to annotate gene families with defense system function with HMMs; Detection (yellow) - time to detect defense systems; and Projection (red) - time to project defense systems across genomes. The benchmark was conducted on a Linux server with two Intel® Xeon® Gold 6150 processors (36 cores), 376 GiB RAM, running CentOS v7.9.2009 with Python 3.10. Execution time scales with dataset size, with the projection step exhibiting super-linear scaling.

**S1 Tab.** Computational performance of the workflow across different species. Execution time (in minutes:seconds format) for each step of the analysis workflow across five bacterial species with varying dataset sizes. Load: time to load the pangenome data; Annotation: time to annotate gene families with defense system function from HMMs; Detection: time to detect defense systems; Projection: time to project defense systems across genomes. Analyzes were performed on the, RAM disponible, type de machine]. Execution time scales with the number of genomes, with the projection step showing the most substantial increase in larger datasets.

| Species | Number of genomes | Load | Annotation | Detection | Projection |
| --- | --- | --- | --- | --- | --- |
| *C. freundii* | 79 | 01:22.18 | 00:09.07 | 01:32.47 | 01:17.98 |
| *P. aeruginosa* | 941 | 02:33.70 | 00:20.60 | 03:47.50 | 08:22.23 |
| *S. enterica* | 1380 | 15:38.76 | 00:12.10 | 13:06.14 | 13:09.22 |
| *K. pneumoniae* | 1659 | 18:51.78 | 00:17.69 | 18:15.45 | 21:04.96 |
| *E. coli* | 3083 | 21:36.71 | 00:26.32 | 26:26.29 | 188:02.75 |

**S2 Fig.**
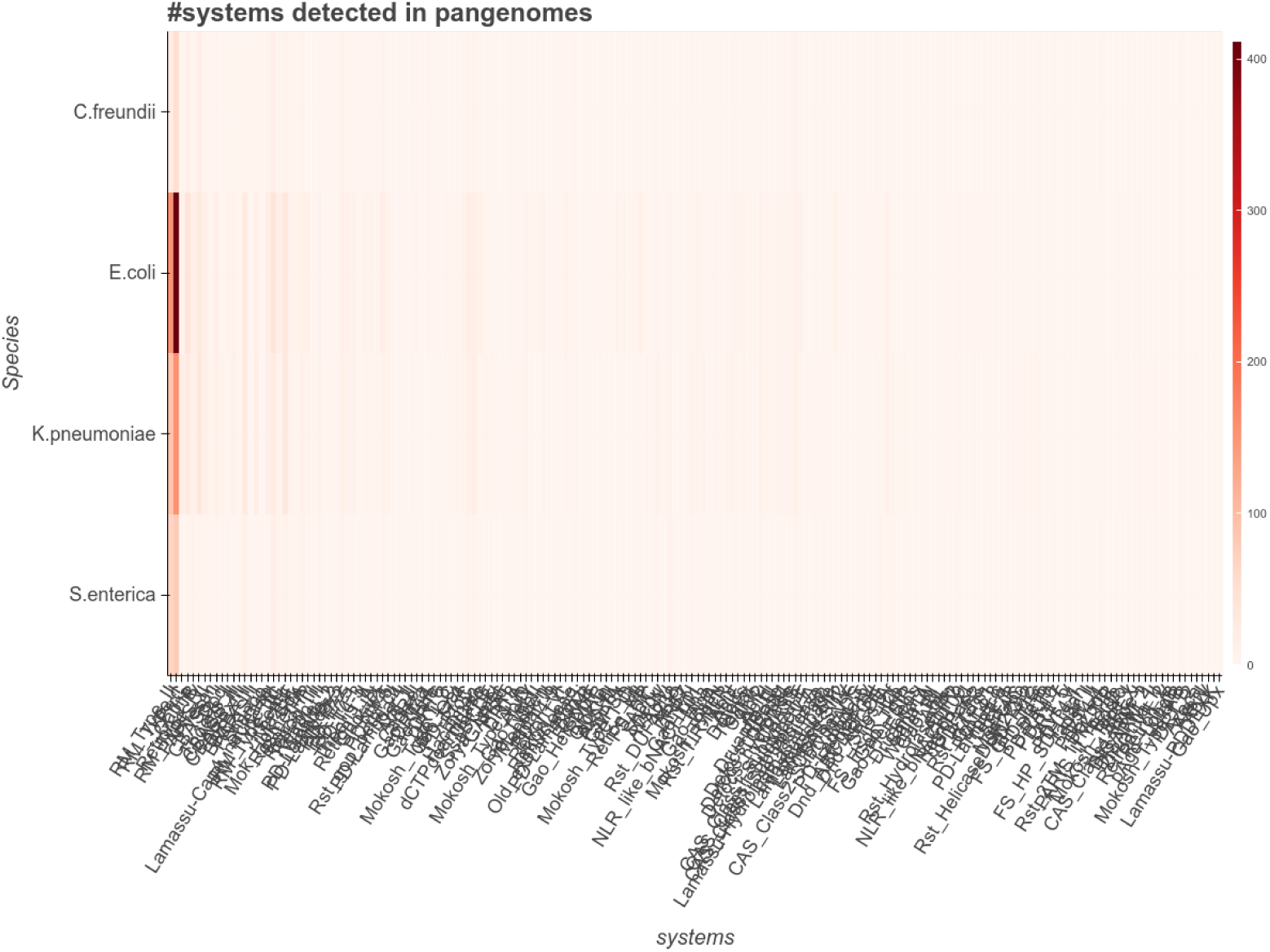
Number of systems predicted for each model in Enterobacteriaceae pangenomes. This figure is automatically generated by PANORAMA. PANORAMA employs the Bokeh package [38] to create interactive and dynamic visualizations, which can be directly accessed and manipulated in any standard web browser.

**S3 Fig.**
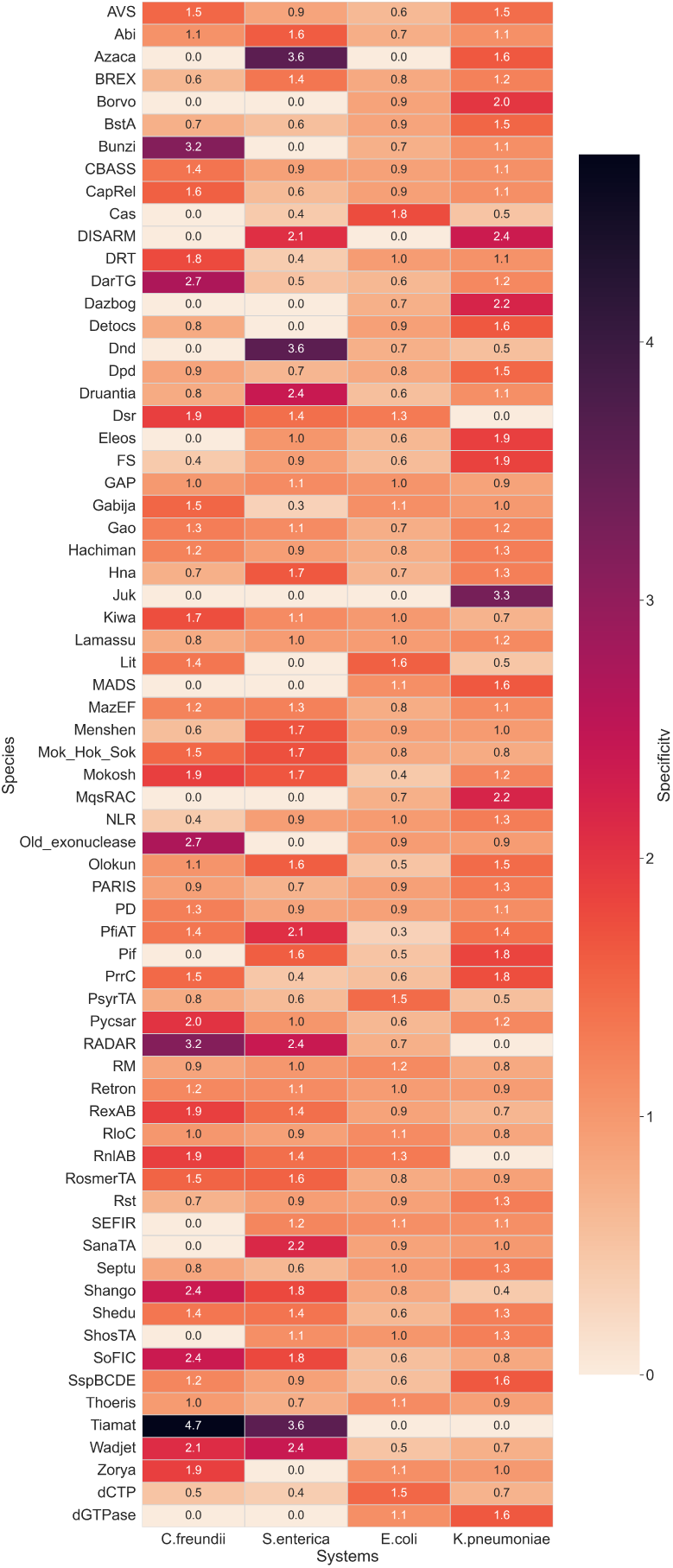
Species-specificity evaluation across system categories in Enterobacteriaceae Pangenomes. The enrichment factors were computed using the method described in Equation 6, providing a quantitative measure of species-specific representation.

**S4 Fig.**
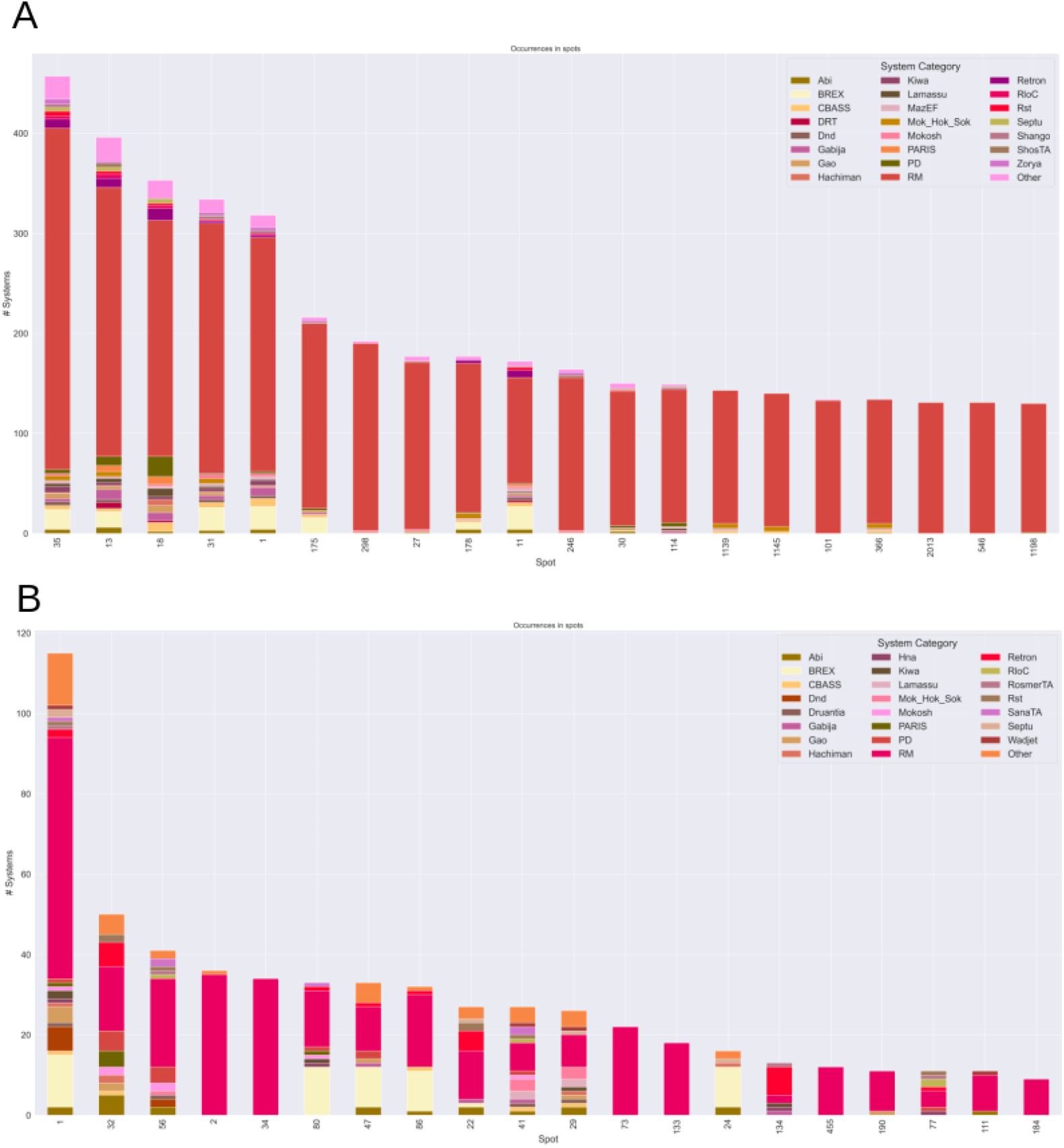
Defense systems within insertion spots. The bar plot displays the number of predicted defense systems in the pangenome for each spot of insertion. (A) *E. coli* pangenome. (B) *S. enterica* pangenome.

**S5 Fig.**
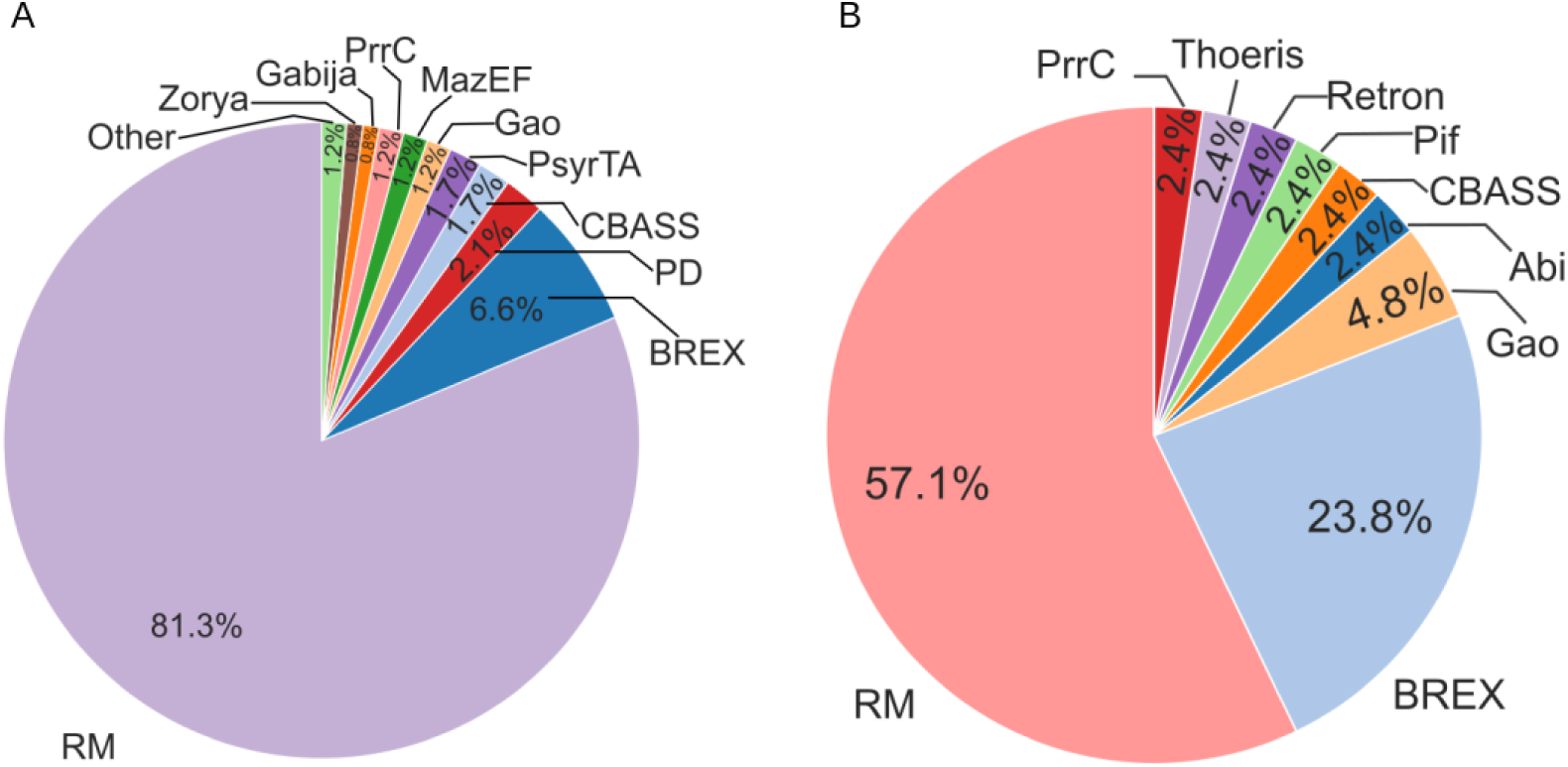
Defense system composition of spots belonging to cluster 3062. Distribution of defense systems in two representative spots from cluster 3062: spot 175 in *E. coli* (A) and spot 86 in *S. enterica* (B). Each slice represents the proportion of a given defense system category within the spot. RM (Restriction-Modification) systems dominate both spots, accounting for 81.3% in *E. coli* and 57.1% in *S. enterica*. BREX represents the second most abundant system in *S. enterica* (23.8%), while it is present at lower frequency in *E. coli* (6.6%). Other defense systems are present at lower frequencies.

## Acknowledgments

We are grateful to Nicolas Wiart for technical support and assistance in testing the software on the computing infrastructure, to Sophie Abby for her help and advice on system modeling, and to Violette Da Cunha for insightful discussions that helped refine the methodological choices during tool development.

## Funding

This research was supported in part by the CFR PhD program of the French Alternative Energies and Atomic Energy Commission (CEA) for JA and the BlueRemediomics project for JM, which is funded by the European Union under the Horizon Europe Program, Grant Agreement No. 101082304. The funders had no role in study design, data collection and analysis, decision to publish, or preparation of the manuscript.

## Data Availability

All data and code necessary to reproduce the findings of this study are publicly available without restriction.

## Data

The complete dataset, including annotated pangenomes and intermediate analysis files, is available from Zenodo [39] at 10.5281/zenodo.15551255. Raw genomic sequences were obtained from RefSeq; the list of RefSeq accession numbers for all genomes used in this study is provided in the Zenodo repository. Commands to download these genomes programmatically are included in the reproduction notebook.

## Code for analysis and reproducibility

PANORAMA (version 1.0.0) used in this study is available on GitHub at https://github.com/labgem/PANORAMA/tree/1.0.0 under the CeCILL v2.1 license and is archived on Zenodo at 10.5281/zenodo.17877647.

A repository containing a Google Colaboratory notebook with all the code to reproduce the figures and analyzes presented in this manuscript, including automated downloading of genomic data from RefSeq, is available on GitHub at https://github.com/labgem/PANORAMA_article and is archived on Zenodo at 10.5281/zenodo.17877540. The notebook can be run directly in Google Colaboratory at https://colab.research.google.com/github/labgem/PANORAMA_article/blob/ main/main.ipynb.

## Author Contributions

JA: Conceptualization, Methodology, Software, Data curation, Investigation, Validation, Formal Analysis, Visualization, Writing – original draft, Writing – review & editing

JM: Methodology, Software, Writing – review & editing

LB: Methodology, Software, Investigation, Writing – review & editing

QFDG: Methodology, Software, Investigation, Validation, Writing – review & editing

YH: Methodology, Software, Investigation, Validation, Writing – review & editing

AC: Conceptualization, Supervision, Funding acquisition, Project administration, Writing – review & editing

DV: Conceptualization, Supervision, Funding acquisition, Project administration, Writing – review & editing

## Notes

### Competing Interest Statement

The authors have declared no competing interest.

